# The origin and evolution of pollen transport in bees (Hymenoptera: Anthophila)

**DOI:** 10.1101/2021.09.19.460919

**Authors:** Zachary M. Portman

**Affiliations:** Department of Entomology, University of Minnesota

**Keywords:** Apoidea, pollen gathering, pollen feeding, *Perdita*, Masarinae

## Abstract

The ability to transport pollen from flowers back to the nest represents a key innovation in the evolution of bees from predatory wasp ancestors. Currently, the origin and evolution of pollen transport remains unsettled. Older hypotheses proposed that crop transport was the original mode of pollen transport, but more recent molecular phylogenies have cast doubt on that view. Instead, more recent hypotheses contend that external transport of dry pollen is ancestral in bees. Here, I propose a new hypothesis to explain the origin and subsequent evolution of pollen transport in bees. I propose that pollen transport arose from adult pollen-feeding behavior and that internal transport of pollen is ancestral in bees. This then led to the evolution of external moist transport, which first required a transition step whereby pollen is temporarily accumulated on the venter on a patch of specialized hairs. Finally, external glazed and dry transport evolved from external moist pollen transport, and the evolution of dry transport led to changes in the location of scopae from the original location on the hind tibia and basitarsus. I illustrate many of these hypothetical evolutionary steps using modern-day bee behavior as an example, with a particular focus on the bee *Perdita tortifoliae*. Examination of the evolution of pollen transport of pollen wasps (subfamily Masarinae) reveals that they have undergone a parallel evolutionary change. Overall, I lay out a broad hypothetical framework to explain the origin and subsequent evolution of pollen transport in bees. This marks a return to the earlier hypothesis that crop transport is ancestral, and it also represents the first in-depth hypothesis to explain how external transport of moistened pollen could have evolved. The evolutionary history of bees has many implications for the biology of bees in the present day, and I lay out a number of predictions that could help confirm or refute my hypotheses.

## Introduction

The evolution of bees from predatory wasp ancestors is one of the major evolutionary developments within Hymenoptera. The shift from a predatory to vegetarian lifestyle may explain the relatively rapid diversification of bees compared to their closest relatives (Branstetter et al. 2017, Murray et al. 2018). In addition, the origin of bees approximately 120 mya coincides with the diversification of early flowering plants (Cardinal and Danforth 2013). However, the mechanisms of how bees originally evolved from predatory wasps remains shrouded in mystery. In particular, the shift from hunting prey to gathering and transporting pollen would require major changes in both morphology and behavior.

The lack of information on the biology of close relatives of bees, combined with the lack of fossils of early bee lineages, make reconstructing the genesis and evolution of bees difficult (Engel 2001, Michez et al. 2012). Recent molecular phylogenies of Hymenoptera suggest pemphredonine wasps are the sister group to bees (Debevec et al. 2012, Branstetter et al. 2017, Peters et al. 2017, Zheng et al. 2018). Sann et al. (2018, 2021) pointed to the Ammoplanidae (also known as the pemphredonine subtribe Ammoplanina) as the closest relatives of bees. The biology of relatively few species of Ammoplanidae are known. These wasps are often found on flowers and the few species for which the biology is known provision nests with Thysanoptera (Maneval 1939, Bohart and Grissell 1972). Other related groups provision with Hemiptera, Collembola, or Thysanoptera (Bohart and Menke 1976). Since the prey preferences of such a small number of species are not necessarily representative of all the species in the group, this tells us little information other than the ancestor of bees potentially visited flowers to find prey and needed to use many prey items (and thus provisioning trips) to complete their provisions (Malyshev 1969).

One of the most important unanswered questions regarding the genesis of bees is how they evolved to transport pollen from flowers back to the nest. In the present day, there exist multiple modes of pollen transport. Bees can transport pollen either internally (in the crop) or externally on specialized structures composed of specialized hair brushes (scopae) or flattened plates (corbiculae) (Thorp 1979). External pollen transport can be further broken down into three modes: pollen can be transported completely dry, completely moist, or glazed, where moist pollen is packed on top of dry pollen (Portman and Tepedino 2017). The evolutionary sequence between the different modes of pollen transport is currently unsettled and it is not clear why multiple modes of pollen transport exist.

The traditional hypothesis has been that the original bees transported pollen internally in the crop (Müller 1883, Malyshev 1936, 1969, Michener 1965, 1979, Jander 1976, Lanham 1980). However, that hypothesis was based in part on the idea that the family Colletidae was basal due to its short, bilobed glossa, a character shared with many Sphecid wasps (Michener 2007). Because many Colletidae transport pollen in the crop (e.g. Euryglossinae and Hylaeinae), this offered a simple and straightforward solution to the problem of how ancestral, wasp-like bees transported pollen (Michener 1979). However, advances in bee phylogenetics have supported Melittidae, rather than Colletidae, as the basal bee family (Danforth et al. 2006, 2012). The family Melittidae contains no known species that transport pollen internally, causing the crop-transport hypothesis to fall out of favor (Michener 2000, Danforth et al. 2019).

More recently, the favored hypothesis is that the original bee transported pollen dry on external scopal hairs (Michener 1944, 2000, 2007, Roberts and Vallespir 1978, Radchenko and Pesenko 1996, Westerkamp 1996, Engel 2001). The more detailed explanations of this hypothesis propose that the protobee, “the hypothetical most recent common ancestor of all bees” (Michener 2000), carried pollen on unspecialized hairs over most of the body surface, and then over time the generalized body hairs specialized and coalesced into discrete structures (Radchenko and Pesenko 1996, Michener 2007). For example, in some bees they formed scopa on the abdominal venter (the family Megachilidae) while others formed scopa on the hind legs (the families Andrenidae and Halictidae). However, one of the main problems with this hypothesis is that the wasp ancestors of the original bee likely did not have copious body hairs, as the closest wasp relatives to bees, the pemphredonine wasps, are small and largely hairless. More recently, Sann et al. (2018), pointing to the evolutionary relationship with pemphredonine wasps, proposed that the ancestor of bees could have transitioned to pollen provisioning by carrying pollen-covered thrips. However, they did not propose any actual mechanism to explain how that could have led to scopae and the deliberate transport of pollen.

Recently, Portman and Tepedino (2017) questioned the hypothesis that external dry transport is ancestral. This was based on an examination of the patterns of evolution of pollen transport in the genera *Perdita* (Andrenidae) and *Hesperapis* (Melittidae); in both genera, it was found that moist pollen transport was the most likely ancestral state and glazed or dry transport were the derived states. This raised the intriguing possibility that external moist transport represents the ancestral state of bees as a whole (Portman and Tepedino 2017). However, we did not propose a potential mechanism for how this could occur, and to date, no studies have proposed an explanation for how moist transport could have evolved, regardless of whether it represents the original pollen transport mode or evolved from another existing pollen transport mode such as dry transport. The only hypothesis I am aware of that even touches on it is by Michener et al. (1978), which suggests the corbiculae in Apidae arose from ancestral “brushy” scopae on the hind legs, potentially as a way to transport sticky nest materials.

Finally, the evolution of the protobee can be informed by the parallel evolution of wasps in the vespid subfamily Masarinae, which have also evolved to provision their larvae with pollen. Bees and pollen wasps both arose around a similar time in the mid-Cretaceous (Branstetter et al. 2017, Peters et al. 2017). All known masarid wasps transport pollen in the crop, making it unambiguously the ancestral trait. Further, exploring the differences in their evolution can help explain why bees are so much more diverse than masarid wasps, with approximately 20,000 species in bees (Michener 2007) vs. approximately 300 species in Masarinae (Carpenter 2001).

The purpose of this paper is to address two specific questions: (1) which mode of pollen transport is ancestral in bees? And, (2) how did the ancestral state of pollen transport diversify into the different modes (internal, moist, dry, glazed) seen in the present day? To address these questions, I follow two primary lines of evidence. First, I use present day bee behaviors (specifically pollen transport, pollen gathering, and pollen feeding) to construct hypotheses regarding how pollen transport originated and transitioned from one mode to another. Second, I examine the biology of masarid pollen wasps, which have undergone a parallel transition to pollen provisioning from predatory ancestors. This approach follows the strategies used by Malyshev (1969) and Jander (1976), but my investigation benefits from recent advances in bee phylogenetics and the greatly increased knowledge of apoid and masarid biology.

Unexpectedly, my conclusions match those of Malyshev (1969) and Jander (1976) in supporting crop transport as ancestral in bees. I further propose that external transport of moistened pollen evolved from crop transport, and I propose a sequence of steps that could result in that transition. The evolution of external moist transport from crop transport is supported by three primary lines of evidence. First, the behavioral steps involved in moistening pollen for transport involve extraneous steps that appear to represent evolutionary vestiges. Second, the similarity of the behavioral steps involved in eating pollen and moistening pollen suggest a shared evolutionary origin. Third, I examine parallel patterns of evolution that have occurred in masarid wasps that may represent transitional evolutionary steps that occurred in bees. Finally, I propose that external dry transport evolved from moist pollen transport and that this led to the expansion and migration of scopal hairs in many bee lineages.

## Methods

Observations of bees took place primarily in Utah and Nevada. *Perdita tortifoliae* Cockerell was observed in the vicinity of St. George Utah, in 2016 and 2017. *Macrotera latior* (Cockerell) and *Hesperapis “timberlakei”* (manuscript name from Stage (1966)) were observed in April 2017, in Lake Mead National Recreation Area. Identifications were made with reference the following taxonomic resources: *Perdita tortifoliae:* Timberlake (1968) and comparison to specimens in the Bee Biology Systematics Laboratory (BBSL) collection; *Macrotera latior:* Danforth (1996) and comparison to specimens in the BBSL collection; *Hesperapis “timberlakei”* MS name: Stage (1966) and comparison to specimens in the BBSL collection. Representative specimens were collected and are deposited in the BBSL collection. Collections of bees in Lake Mead National Recreation Area were made under permit #LAKE-2017-SCI-0004.

A Quanta FEG 650 Scanning Electron Microscope was used to image the specimen hairs and videos were taken with a Sony A65 DSLR camera and edited using Sony Movie Studio 13 software.

## Results and Discussion

### The roadmap

The basic steps in the origin and evolution of pollen transport follow the general sequence of crop transport -> external moist transport -> external dry transport.

1. Crop transport represents the original form of pollen transport and evolved from pollen feeding behavior. Bees consumed pollen by nibbling with the mouthparts and by drawing a pollen-covered foreleg through the mouthparts.
2. The next stage of pollen transport evolution was the accumulation of pollen on the venter. The accumulated pollen was then picked up by the foreleg and brought forward to the mouthparts and consumed by drawing the foreleg through the mouthparts.
3. Next, external moist pollen transport evolved from internal transport, likely due to leftover pollen becoming stuck to the hind leg rather than completely groomed off.
4. External dry and glazed transport evolved from external moist transport in parallel with a development of the scopal hairs, following the hypothesis of Portman and Tepedino (2017).
5. Finally, in various lineages that transport dry pollen, the scopal hairs expanded and migrated from the hind tibia and basitarsus towards the midline of the body. In other lineages, crop transport secondarily evolved.

What follows is a rather meandering discussion of the evidence supporting this roadmap. This is then compared to the hypothesized parallel evolution of pollen transport in pollen wasps.

### Hypothesis: Crop transport is ancestral and it evolved from ancestral adult pollen-feeding behavior

In the present day, pollen feeding is an integral part of bee biology; pollen is eaten by adult bees (both male and female) and is necessary for the production of eggs (Robertson 1929, Rozen 1989, 1958, Stockhammer 1966, Shinn 1967, Jander 1976, Batra 1985, Hunt et al. 1991, Richards 1994, Michener 2007, Schäffler and Dötterl 2011, Cane 2016, Cane et al. 2016, Houston 2019). While gathering pollen, females will often take a bite to eat without interrupting pollen gathering activities (Jander 1976, ZP pers. obs.).

The ubiquity and importance of pollen-feeding in bees suggests a basal origin, and it is simple to hypothesize how ancestral pollen-feeding behavior could evolve into transport of pollen in the crop. In this case, it would require adults of the protobee to first consume pollen and nectar (or other plant exudates) for its own energetic and nutritional needs. Despite the limited fossil record, there is direct evidence that aculeate wasps fed on angiosperm pollen for their own nutritional needs as early as the cretaceous (Grimaldi et al. 2019). The next step in the evolution of pollen transport requires the protobee to regurgitate the consumed pollen and nectar back at the nest. The specific behaviors and mechanisms by which regurgitation evolved are unknown. However, regurgitation of food to provision the young has evolved multiple times in multiple different Hymenopteran lineages including ants, pollen wasps and other vespids (Liebig et al. 1997).

This hypothesis — that crop transport evolved from pollen feeding — has been previously proposed by Malyshev (1969) and Jander (1976). The strongest argument against it is that there are no known examples of basal bees that transport pollen in the crop. However, there are two points that support this hypothesis: first, essentially all bees that have had their biology explored in depth feed on pollen and regurgitate nectar onto their larval provisions. These two behaviors may represent evolutionary vestiges of ancestral crop transport. Second, as I will explain subsequently, crop transport provides transition steps that are necessary for the next stage of pollen transport: the evolution of external moist transport.

### A discussion of the mechanisms by which bees feed on pollen

In order to understand the evolution of pollen transport, it is first necessary to have a thorough understanding of the specific steps bees use to feed on pollen. All known bees consume pollen by drawing the foreleg through the mouthparts (Jander 1976, Michener 2007). Use of the foreleg for consuming pollen represents a modification of typical Hymenopteran grooming behavior. In most other Hymenoptera, the foreleg is cleaned by drawing it through the mouthparts (Farish 1972, Jander 1976). However, in bees, this movement has been co-opted for pollen feeding — indeed, the majority of bee groups have a comb on either the galea or stipes that is specifically used for scraping pollen from the foreleg (Jander 1976). Supporting this hypothesis that ancestral foreleg grooming has been co-opted for pollen feeding is the fact that bees are potentially unique among Hymenoptera in grooming the foreleg by pulling it through the bent midleg (Farish 1972, Jander 1976; see Fig. 3 of Jander 1976 for illustration). In other words, the ancestral method of foreleg-cleaning (drawing it through the mouthparts) has been replaced by a derived method of foreleg-cleaning (drawing it through the crook of the midleg).

The use of the foreleg for consuming pollen presents a puzzle since presumably the simplest way to consume pollen would be to nibble it directly with the mouthparts. Indeed, bees are capable of nibbling pollen directly with the mandibles and have been observed to do so when consuming pollen directly from pollen masses in the nest (e.g. Batra 1964), but they apparently do not perform this behavior on flowers (Jander 1976). This is likely because using the forelegs for pollen consumption offers two main advantages: first, it allows consumption of pollen from any place the foreleg can groom, namely the head and thorax (Jander 1976). This allows bees to exploit pollen that has been deposited on the head or thorax by a flower. Second, nibbling pollen presents mechanical difficulty in that pollen is difficult to swallow. In order to be easily swallowed in a large quantity, pollen must be mixed with regurgitated nectar (or potentially some other fluid), a process I have frequently observed performed by bees eating pollen, and pollen feeding behavior is often performed in tandem with nectar concentrating behavior (see Portman et al. In Press). The hypothesis that feeding on pollen requires a liquid such as nectar is supported by the fact that bees dissected after eating pollen have a mix of pollen and nectar in the crop (Danforth 1989, 1990, Cane et al. 2016). Other pollen-feeding Hymenoptera face this same problem but solve it by a different route: Muttilidae and Scoliidae regurgitate liquid directly onto anthers before consuming the pollen (Jervis 1998).

### A window back in time: Modern day pollen-feeding behavior is essentially the same as ancestral pollen transport behavior

Eating pollen for adult nutrition and consuming pollen in order to transport it back to the nest are functionally equivalent behaviors. The only real difference is whether or not the bee regurgitates the pollen back at the brood cell (provisioning) or digests it (feeding). As a result, a careful study of the mechanisms by which bees feed on pollen can provide a template for how the protobee transported pollen.

Here, I use the bee *Perdita tortifoliae* as the archetypal bee to demonstrate pollen feeding and pollen gathering behavior. I use this bee primarily because I have been able to make a close and careful study of its pollen-feeding and pollen-gathering habits. *Perdita tortifoliae* is a minute bee (about 4 mm body length) that specializes on the pollen of *Lepidium* (Brassicaceae), which it transports moistened on the hind legs. It occurs in the arid western United States and is locally common in the vicinity of St. George Utah, where observations took place.

### Pollen-feeding behavior in *Perdita tortifoliae*

Similar to other bees, the females of *P. tortifoliae* will occasionally take bites of pollen while gathering pollen and packing it into their scopae. However, towards the end of their daily activity on flowers in the afternoon, *P. tortifoliae* females engage in dedicated feeding trips where they exclusively consume pollen without packing any into the scopa. Indeed, any excess pollen is totally discarded. This feeding trip is presumably the same as feeding trips in other panurgine bees, who return to the nest with empty scopa but have pollen and nectar in the crop (Danforth 1989, Neff and Danforth 1991, Visscher and Danforth 1993). These previously observed feeding trips have only been observed through dissecting bees returning to their nests and, to the best of my knowledge, this behavior of feeding on pollen in panurgines has not previously been reported. One of the most important features of the pollen-feeding behavior of *Perdita tortifoliae* is that the bees first accumulate pollen on a specialized patch of hairs on the venter (as in Fig. 4A–B), a strategy that is well-documented in the pollen-gathering behavior of various panurgine bees (Portman et al. 2019).

The behavior of pollen feeding can be divided up into five main steps (see also Figure 1 and Supplemental Video 1: https://youtu.be/6M4BpnQ8zfc):

**Accumulating pollen:** The forelegs (and occasionally the midlegs) are used to scrape pollen directly from anthers and deposit it on the venter of the thorax.
**Unloading pollen:** After a sufficient quantity of pollen has accumulated on the venter of the thorax, the bee rears back on its hind legs, often forming a tripod with the apex of the abdomen. Pollen is removed from the venter by the forelegs using from one to ten downward scraping motions.
**Bringing pollen forward:** The legs with pollen are brought to the mouthparts and the tongue is extended, and the bee regurgitates nectar onto the base of the mouthparts.
**Eating the pollen:** One at a time, each foreleg is drawn through the mouthparts, either in between the split galeae or between the closed galeae and a mandible. Steps 2–4 are then repeated until the pollen has been removed from the venter of the bee.
**Discarding excess pollen:** During the whole process, excess pollen is continuously groomed off of the front legs by scraping them through the crook formed by the inner side of the mid-femur and mid-tibia, and the midlegs are in turn scraped through the crook of the hindlegs. The pollen is then groomed and discarded by the hindlegs rubbing against each other.

**Figure 1.**
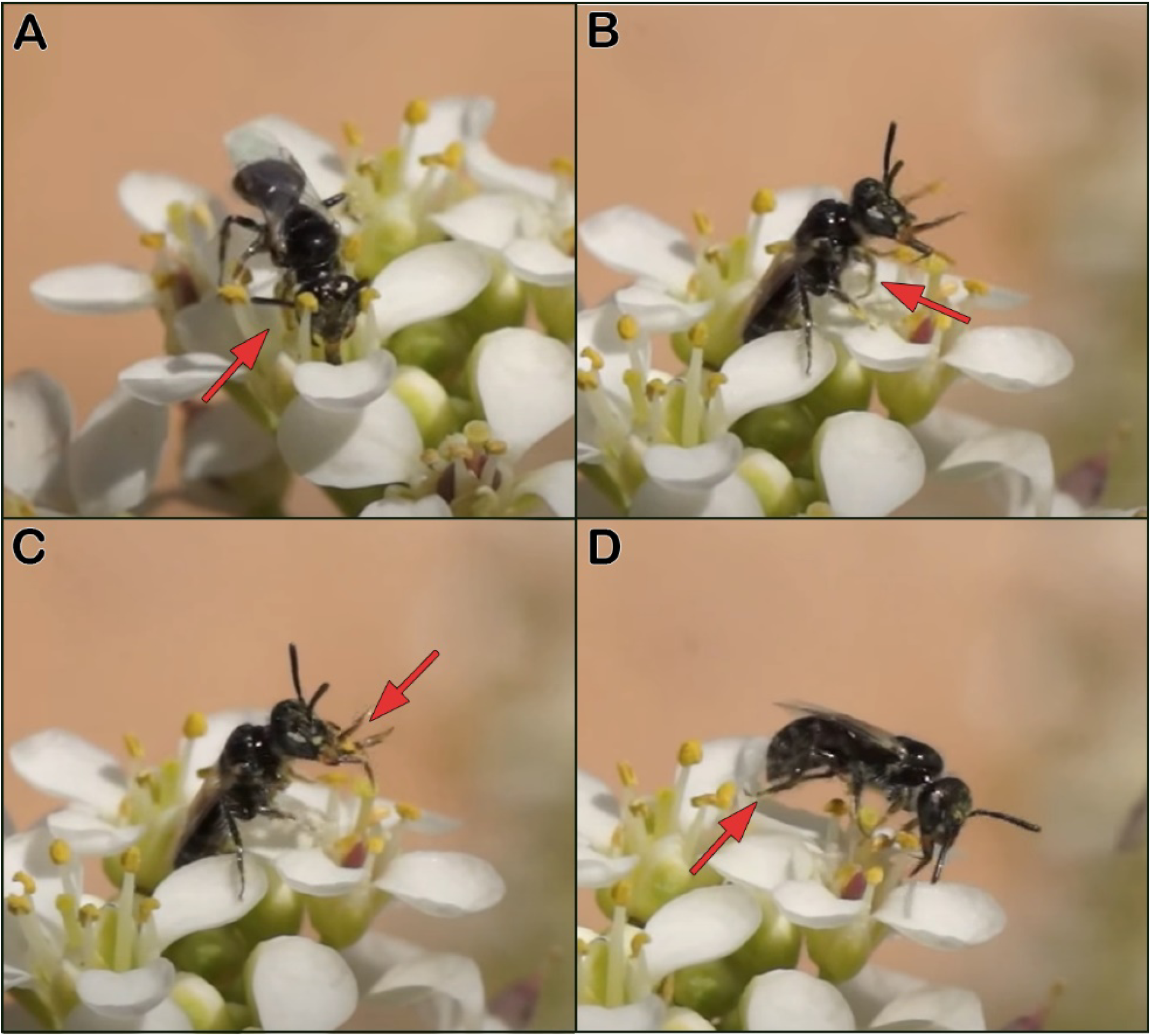
Pollen-feeding behavior in *Perdita tortifoliae* on *Lepidium* flowers. A) Using the foreleg to accumulate pollen on the venter B) Rearing back and using the forelegs to remove the pollen from the venter. C) Splitting apart the mouthpart and drawing the pollen-covered foreleg through the mouthparts D) Using the hind legs to remove excess pollen. It is much clearer in the video, available as Supplemental video 1: https://youtu.be/6M4BpnQ8zfc

#### Hypothesis: this represents the ancestral form of pollen transport

I propose that this mode of feeding on pollen represents the ancestral form of gathering pollen. It is a key point that the pollen is accumulated on the venter prior to being consumed. It seems probably that the accumulation of pollen on the venter step evolved after pollen transport in the crop, though it could have evolved before crop transport as a way to more efficiently feed on pollen. However, regardless of when it evolved, it is a necessary preadaptation to evolve external pollen transport. This will be made clear by a comparison of pollen-feeding and external pollen transport in *Perdita tortifoliae* in the next section.

### The evolution of external moist pollen transport from pollen feeding behavior

One of the key points in my argument is that only a couple minor changes are required to turn pollen feeding into external pollen transport. I will demonstrate this here by describing pollen-gathering and packing behavior of *Perdita tortifoliae* and comparing it to pollen-feeding behavior in *P. tortifoliae* that was described in the previous section. I then show how pollen-feeding can evolve into external moist pollen transport with just some minor changes.

#### Pollen gathering and packing behavior in *Perdita tortifoliae*

Like other panurgines, and similar to how it feeds on pollen, *P. tortifoliae* gathers pollen using a two-step process, where it temporarily accumulates pollen on a specialized patch of apically hooked hairs on the venter of the thorax before transferring it to the hind legs (as in Fig. 4A–B; reviewed in Portman et al. 2019). It also moistens the pollen before packing it onto sparse scopae for transport (Portman and Tepedino 2017). Here, I further break it down into finer steps in order to better illustrate the component behaviors (see also Figure 2 and Supplemental video 2: https://youtu.be/v1G96DLynCQ).

**Accumulating pollen:** The forelegs (and occasionally the midlegs) are used to scrape pollen directly from anthers and deposit it on the venter of the thorax.
**Unloading pollen:** After a sufficient quantity of pollen has accumulated on the venter of the thorax, the bee rears back on its hind legs, often forming a tripod with the apex of the abdomen. Pollen is removed from the venter by the forelegs using from one to ten downward scraping motions.
**Bringing pollen forward:** The legs with pollen are brought to the mouthparts, the tongue is extended, and the bee regurgitates nectar onto the base of the mouthparts.
**Moistening the pollen:** Both forelegs are brought up together and scraped across the top of the extended mouthparts, moving from the base to the apex of the mouthparts and picking up regurgitated nectar in the process.
**Transferring and packing the pollen back on the hind legs:** Immediately following pollen moistening, the foreleg is drawn through the midleg, in a crook formed by the inner side of the mid-femur and mid-tibia, causing the pollen to be transferred to the hind part of the midleg, and the midleg then pats back against the hind tibia, depositing the pollen. Steps 3–5 are then repeated until the pollen has been removed from the venter of the bee.

**Figure 2.**
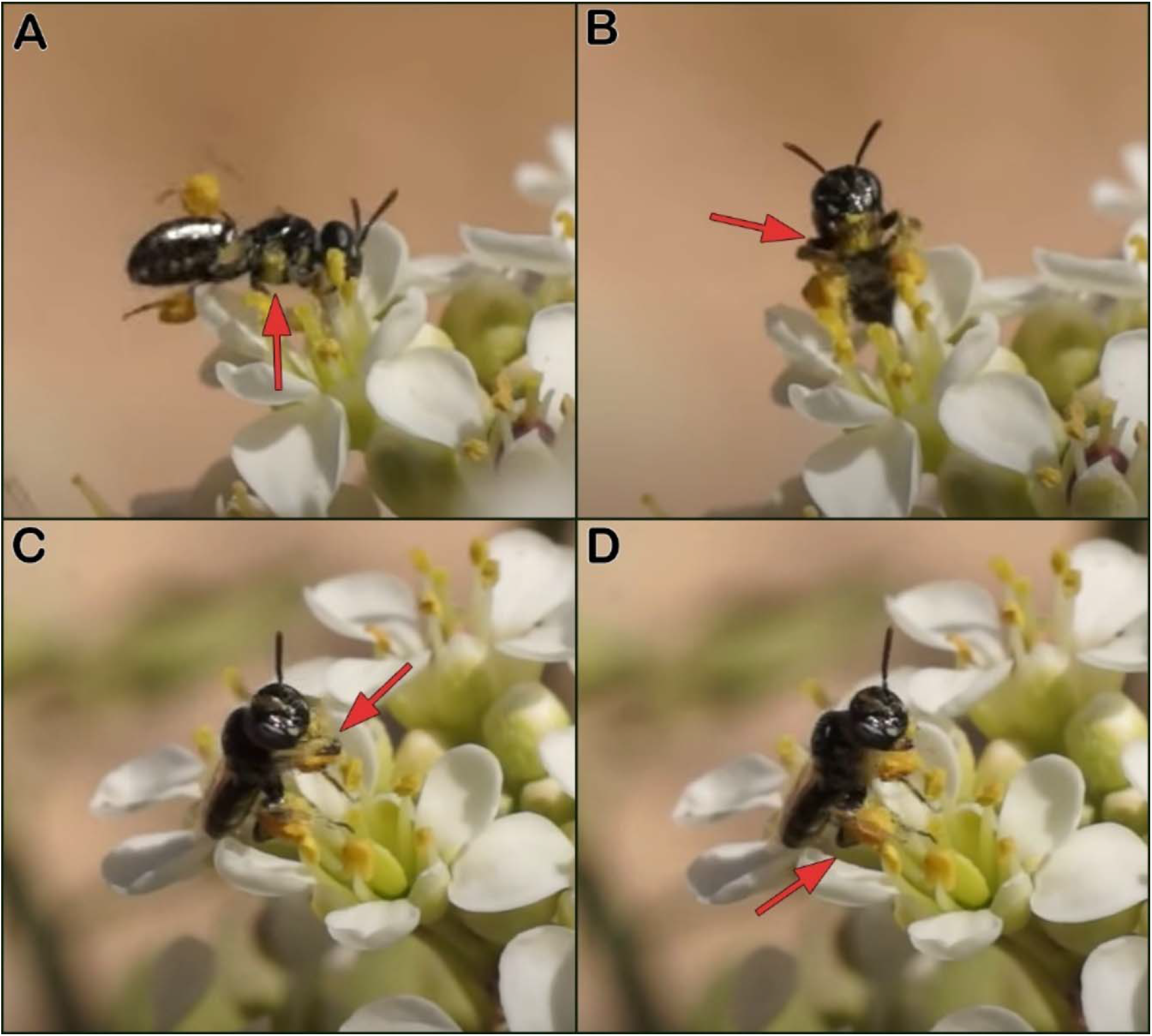
Pollen gathering behavior in *Perdita tortifoliae.* A) Accumulating pollen on the venter. Pollen is visible because the female was just knocked by a male. B) Rearing back and removing the pollen from the venter with the forelegs. C) Drawing the pollen-covered forelegs along the extended mouthparts to moisten them with pollen. D) Using the midlegs to transfer the moistened pollen to the hind legs and tamp it down. It is much clearer in the video, available as Supplemental video 2: https://youtu.be/v1G96DLynCQ

#### Changing to external moist transport from pollen feeding

Comparing pollen feeding to pollen packing behavior in *Perdita tortifoliae* (Table 1), two key points are apparent. First, they represent variations on the same basic behavior, and the first three steps are shared between them. Second, only two changes are needed to go from pollen-feeding to pollen-packing: in pollen-packing, the pollen is brought along the mouthparts to be moistened (rather than consumed) and the pollen is packed onto the hind legs (rather than groomed off). As we have seen, when the bee feeds on pollen, a portion of pollen is already passed along the tongue without being consumed, so packing rather than discarding pollen is the primary step that needs to change.

**Table 1.**
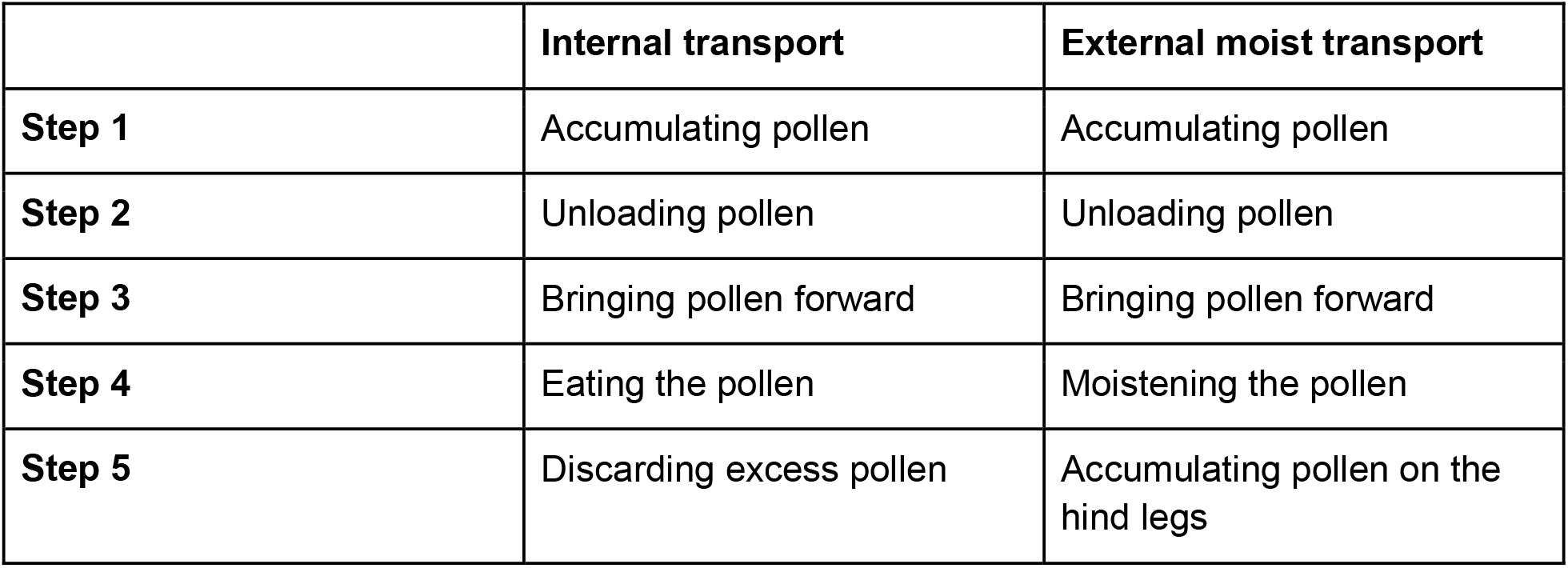
Comparing the steps of internal transport (pollen feeding) behavior vs external moist transport behavior

|  | Internal transport | External moist transport |
| --- | --- | --- |
| **Step 1** | Accumulating pollen | Accumulating pollen |
| **Step 2** | Unloading pollen | Unloading pollen |
| **Step 3** | Bringing pollen forward | Bringing pollen forward |
| **Step 4** | Eating the pollen | Moistening the pollen |
| **Step 5** | Discarding excess pollen | Accumulating pollen on the hind legs |

#### Evolutionary implications

Viewing this from an evolutionary standpoint, this provides the map for how external pollen transport evolved; as a modification of crop transport. In essence, external pollen transport represents internal pollen transport with just a couple of modified steps. Both behaviors center around the temporary accumulation of pollen on the venter; the main difference being that in pollen-feeding, the pollen is consumed by the mouthparts and in pollen-gathering the pollen is moistened by the mouthparts and passed back.

The steps between pollen-feeding and external pollen transport require few to no transition steps. One could imagine a gradual change where internal pollen transport in the crop slowly changes over to external moist transport. In this case, some of the excess pollen from eating would be passed back and glommed onto the legs rather than groomed off completely. Because this pollen had been in contact with the nectar at the mouthparts, it would be moistened and sticky. The bee would instead wait until it was back at the nest to completely groom it off. This amount of pollen on the legs would grow over time, making up a greater and greater proportion of the pollen load, until eventually, external moist transport became the predominant or sole method of pollen transport.

Viewing external moist transport as a behavior that has been tacked onto pollen-feeding behavior explains the incongruous step in pollen gathering behavior, where the bee brings the pollen forward to the mouthparts to be moistened. During that process, the bee picks up the pollen from the venter of the thorax, brings the pollen forward to be moistened at the mouthparts, only to immediately pass the pollen backwards towards the hind legs. This stands in contrast to what seems like the more logical method of simply passing the nectar backwards to moisten pollen (as if often seen in honey and bumble bees). However, bringing pollen forward makes sense because that represents the origin of the behavior from when the bee simply ate the pollen that was brought forward rather than moistening it. In this view, passing pollen forward to the mouthparts to be moistened represents a vestige of the ancestral pollen feeding behavior and this step is retained due to its evolutionary history rather than any particular utility.

If moist transport evolved from internal transport, it explains how external pollen transport could have evolved without any specialized pollen-carrying structures, as *Perdita* (and many other bees that transport moistened pollen) carry pollen on short, sparse, and simple scopal hairs. Indeed, the protobee may have been similar in many ways to *Perdita tortifoliae:* small, relatively hairless, and lacking specialized pollen transporting hairs. The one exception to the lack of specialized hairs is the specialized patch on the venter. Though I do not go into it here, the temporary accumulation of pollen on a specialized patch of hairs (e.g. Fig. 4) provides a potential mechanism for bees to specialize on the morphological properties of pollen despite the lack of specialization in the scopal hairs; this is important given that pollen specialization is increasingly viewed as the ancestral state in bees (Michez et al. 2008, Sedivy et al. 2008).

There are still many unknowns about the exact behavior and evolutionary history of the first bees. It is worth noting the possibility that crop transport is not ancestral, and instead external pollen transport evolved directly from adult pollen-feeding behavior (using the same mechanism just outlined). However, I consider that unlikely, particularly given the parallel evolution of pollen wasps, discussed in a later section. In addition, it seems unlikely that feeding on pollen for adult nutrition would generate enough excess pollen to attach in appreciable quantities on the hind legs, especially in an individual that continues to have a predatory lifestyle. It seems likely that the earliest forms of crop-transporting bees are extinct; given that all known bees — including males and parasitics — share a broadened hind basitarsus (Radchenko and Pasenko 1996, Michener 2007). This suggests that the most recent common ancestor of all extant bees transported moist pollen on the hind legs.

#### Some additional supporting evidence from other bees

While the behavior of *Perdita tortifoliae* was used to illustrate the proposed evolutionary sequence of steps in the evolution of pollen transport, they are by no means a special case. They are merely the ones I had the opportunity to observe the most in-depth, and there are additional bees that have these same behaviors. For example, the same pollen-feeding and pollen-gathering behaviors were observed in the species *Macrotera latior (Macrotera* is the sister genus to *Perdita),* though their faster speed and tendency to transfer the pollen without standing in a tripod position made the behaviors more difficult to observe and record *(M. latior* pollen feeding: Supplemental video 3: https://youtu.be/tdUz_iTr8qY and *M. latior* pollen gathering: Supplemental video 4: https://youtu.be/l6C6KtmqSD8). In addition, the practice of temporarily accumulating pollen on the venter is widespread in other panurgine bees, reported in at least 14 other panurgine species, and it has also been recorded in disparate other groups, including *Trigona* and *Macropis* (reviewed in Portman et al. 2019). Although pollen feeding behavior has not been documented for those species, I see no reason why they would differ from *Perdita tortifoliae* and *Macrotera latior*.

The same pollen feeding and gathering behaviors also occur in the melittid bee *Hesperapis “timberlakei”* Stage (1966) manuscript name (hereafter *H*. *“timberlakei”).* This bee has a preference for *Psorothamnus* pollen but also gathers pollen from *Larrea* (Michez et al. 2008, ZP pers. obs.). It transports moistened pollen on hind leg scopae (Portman and Tepedino 2017). Multiple females of *H*. *“timberlakei”* were observed gathering pollen from *Psorothamnus fremontii,* and a short clip of one was recorded (Fig. 3, Supplemental video 5: https://youtu.be/Tpbd2UrmLls). These observations confirm two key aspects of the pollen gathering behavior of *H*. *“timberlakei”.* First, gathered pollen is initially accumulated on the venter of the thorax by the fore- and midlegs. Second, pollen is passed up to the mouthparts to be moistened before being passed back to the scopae (Fig. 3B). Because the transfer of pollen from the venter took place while the bee was in flight, it was very difficult to observe, though the movements can be discerned when the video is slowed down (Supplemental video 6: https://youtu.be/Wzn37N3sNDc). Investigation of the venter of the thorax of *H*. *“timberlakei”* reveals that, like most *Perdita,* it has a specialized patch of apically hooked hairs where the pollen accumulates (Fig. 4C–D). Finally, in a subsequent review of old videos, I found one of a *Hesperapis* (likely *H. “timberlakei”)* feeding on pollen by first accumulating on the venter, but unfortunately only captured a short and obstructed video (Supplemental video 7: https://youtu.be/NK10JpnzblI).

**Figure 3.**
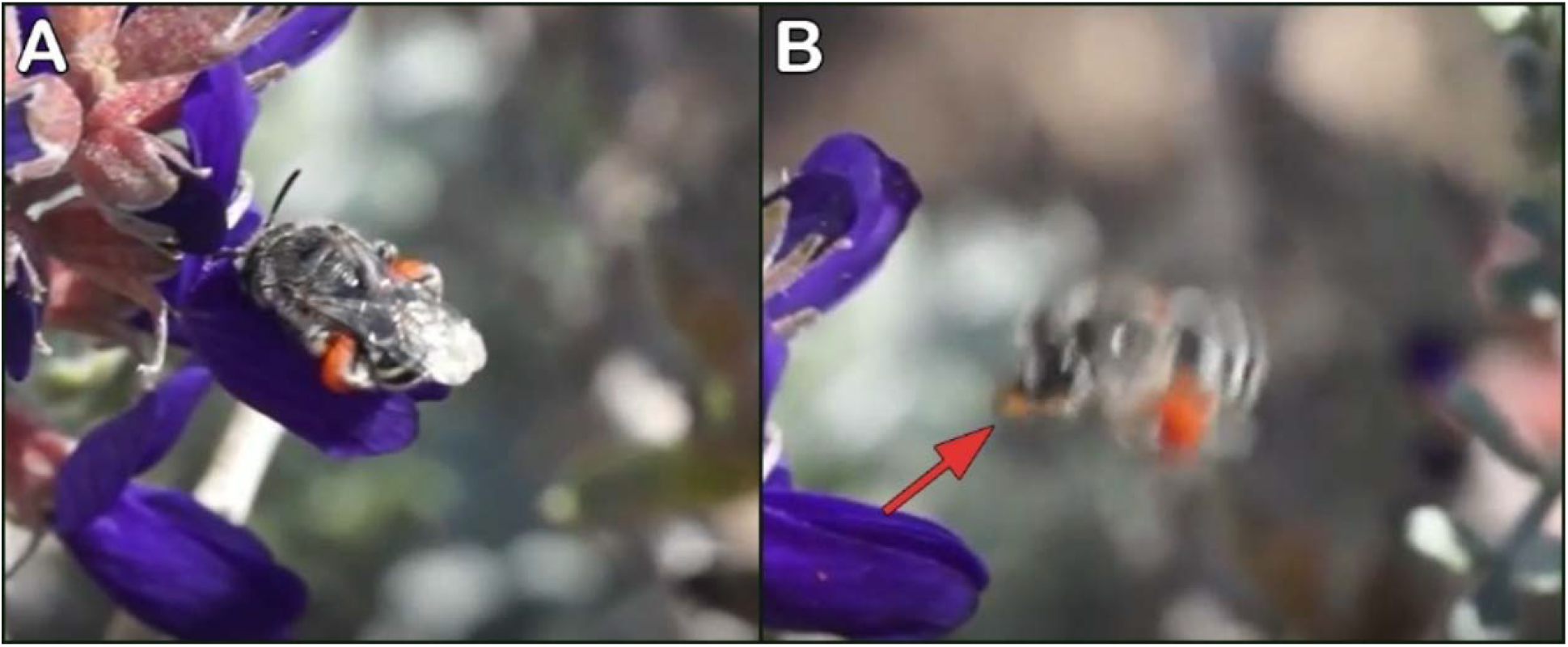
*Hesperapis* “*timberlakei*” MS gathering pollen A) gathering pollen and accumulating pollen on the venter of the thorax. B) Bringing pollen forward to the mouthparts for moistening whilst in flight; red arrow indicating bright orange *Psorothamnus* pollen on the foreleg. See Supplemental video 5: https://youtu.be/Tpbd2UrmLls and Supplemental video 6: https://youtu.be/Wzn37N3sNDc.

**Figure 4.**
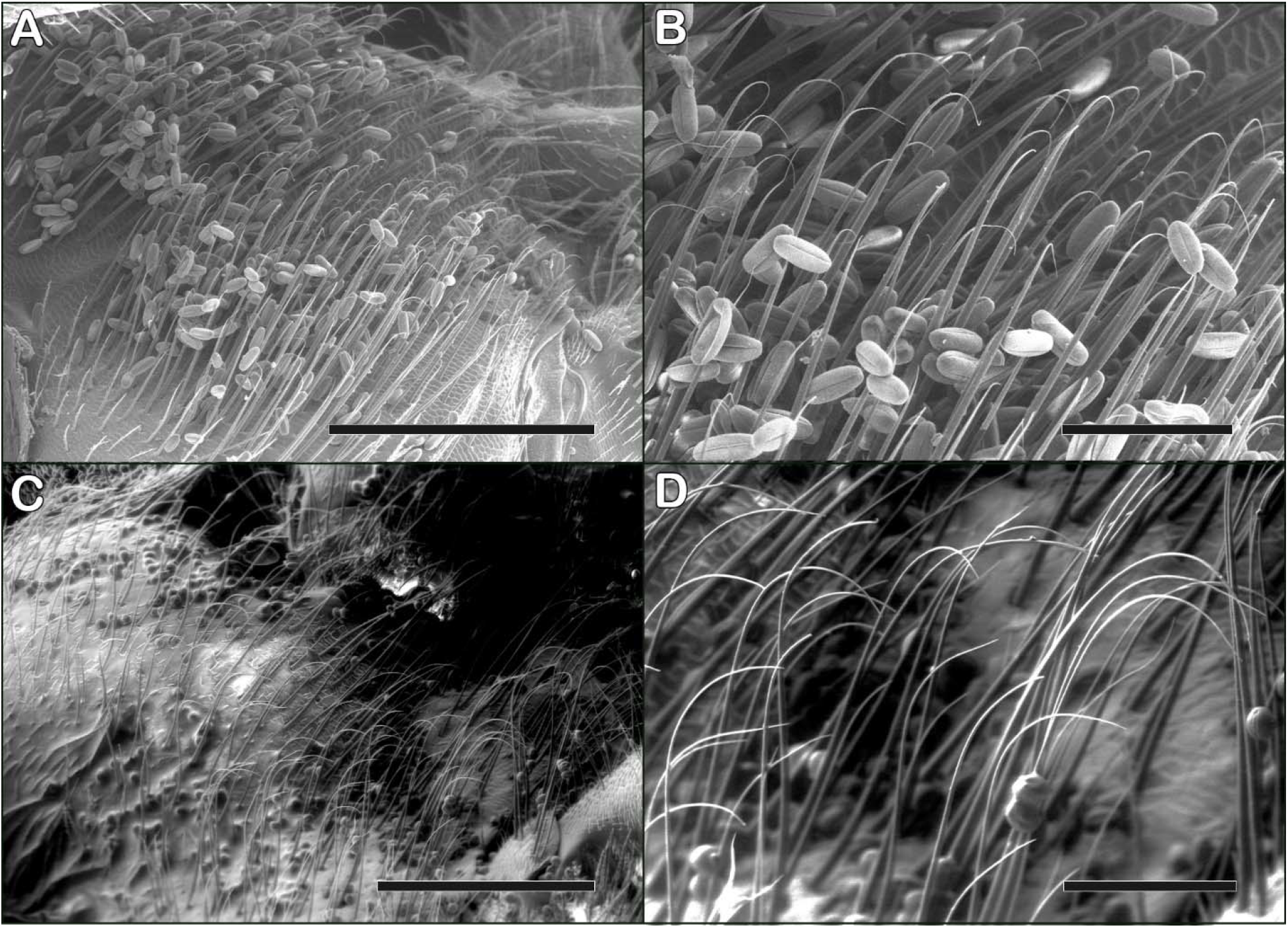
Patch of specialized ventral hairs in *Perdita perpallida* (the ventral patch on *P. tortifoliae* is similar) and *Hesperapis “timberlakei”* MS. Shown is the venter of the thorax; bees are positioned upside-down and with the head to the left. A) *Perdita perpallida,* scale bar = 400 um. B) *Perdita perpallida* scale bar = 100 um. C) *Hesperapis “timberlakei”* MS, scale bar = 400 um. D) *Hesperapis “timberlakei”* MS, scale bar = 100 um.

The similarities between the pollen-gathering and pollen-feeding behavior of *H*. *“timberlakei”* and *P. tortifoliae* are significant because *H*. *“timberlakei”* is in the family Melittidae, the basal bee family. This supports the hypothesis that gathering pollen by first accumulating it on the venter is basal as well. Unfortunately, due to the relative rarity of melittid bees, observations of their pollen gathering and feeding behavior are frustratingly sparse. One other *Hesperapis* species, *H. laticeps,* has been observed temporarily accumulating pollen on the venter, though the pollen is primarily held in genal hair baskets located on the venter of the head rather than the venter of the thorax (Portman et al. 2019). However, other than the location of the ventral hair patch, the pollen gathering movements are similar between *H*. *“timberlakei”* and *H. laticeps,* the and the location of the patch on the head is likely an adaptation to extracting pollen from flowers of *Mentzelia* and *Eucnides.* Other melittid bees in the genus *Macropis,* which transport oil-moistened pollen on hind leg scopae, have been found to also gather pollen by accumulating it on the venter before transferring it to the scopae (Cane et al. 1983, Vogel 1992, Schäffler and Dötterl 2011), suggesting this behavior is likely more widespread but unreported in the family.

#### The evolution of glazed and dry transport from moist pollen transport

The origin of external dry transport likely evolved from moist transport by the stages laid out in Portman and Tepedino (2017). In short, bees that transported moistened pollen underwent an evolutionary transitioned to dry transport by initially packing pollen dry into the scopae before capping it with moistened pollen. This process was facilitated by bees that switched to host plants with adhesive pollen that stayed in the scopae without the need to be agglutinated by nectar. However, due to the short length of the scopal hairs, only a small amount of pollen could be carried dry, and any additional pollen needed to be agglutinated with nectar on top of the initial layer of dry pollen. Over evolutionary time, the proportion of dry pollen gradually increased as the scopal hairs developed and extended and were able to carry greater amounts of dry pollen. One exception occurs in *Perdita* that utilize Onagraceae pollen — as this pollen has naturally occurring sticky viscin threads that are transported most effectively on sparse, simple scopal hairs (Linsley 1958). The end result of this process was that many bees transitioned to completely dry transport, while other species in the present day retain the vestige of this process and still glaze the pollen, or cap it with moistened pollen. One important point from Portman and Tepedino (2017) is that the evolution of dry transport is associated with specialization on certain pollen types, especially spiky or sticky pollen that either makes moist transport less efficient, dry transport easier, or a combination of both.

Glazed pollen transport, where bees initially pack dry pollen into their scopae but then cap it with moistened pollen, appears to be something of a transition state between moist and dry transport (Portman and Tepedino 2017). However, in various species glazed pollen transport appears to be an evolutionary endpoint in and of itself; examples of this include various *Perdita, Hesperapis*, and *Dufourea novaeangliae* (Eickwort et al. 1986, Portman and Tepedino 2017).

No doubt further investigation will reveal more species that transport glazed pollen. It’s unclear why some species continue to glaze pollen rather than evolving entirely dry transport.

One important aspect of the evolution of dry pollen transport is that it often leads to the loss of the transition step where bees temporarily accumulating pollen on a specialized hair patch on the venter. However, this behavior is retained in some bees that transport dry pollen. For example, temporarily accumulating pollen on the venter is retained in *Macrotera* subgenus *Macrotera,* which transports dry Cactaceae pollen in tibial scopae (e.g. Neff and Danforth 1991), while the rest of the genus transports moistened pollen. However, many other lineages that have switched to dry pollen transport lose the temporary accumulating pollen step, and instead directly pass pollen to the scopae, or even gather pollen directly with the scopae by rubbing or tapping the scopae directly against the pollen source as in many Megachilidae (Portman et al. 2019). The loss of the temporary accumulation of pollen in the pollen gathering process makes the evolutionary transition from moist pollen transport to dry pollen transport a one-way street, since that step is generally necessary to transport moistened pollen.

#### The further evolution of dry transport and the shifting of the scopal hairs

The transport of dry pollen is associated with the expansion of the scopal hairs to new areas. All bees that transport moistened pollen transport it exclusively on the hind tibia and basitarsus. The greatest degree of scopal expansion in bees that transport moist pollen is that the pollen carrying area has expanded to the rear of the hind tibia and basitarsus, forming a complete “muff’ of pollen that encircles the leg (e.g. Malyshev 1936, Rozen 1989). In contrast, the transport of dry or glazed pollen is often associated with the expansion of the scopal hairs to entirely new areas of the body. For example, in some species, the transport of glazed (partially dry) pollen is associated with the expansion of the pollen-transporting hairs to more proximal hind leg segments (e.g. Portman and Tepedino 2017). In terms of broad-scale evolutionary trends in bees, there is a parallel change in different bee groups, with the scopal hairs expanding or migrating from the distal to the proximal areas of the body. This is most clearly demonstrated in *Andrena, Colletes,* and various Halictidae, where the majority of pollen is carried on the thorax, sterna, and basal leg segments rather than the hind tibia and basitarsus (Roberts and Vallespir 1978, Michener 1999), which I contend represents the ancestral location of the scopa. Why the scopae have become increasingly proximal is not clear, but it could be an adaptation to better secure the pollen from being scraped off by nesting substrate or forces from wind during flight.

The expansion and migration of scopae is particularly intriguing in the evolution of Megachilidae, which transport pollen on abdominal scopa. For most other bee groups, the migration of the scopae is straightforward, with additional scopal areas being added, but relatively minimal loss of pre-existing scopal structures. For example, in some groups, such as *Colletes* and *Andrena,* the scopal hairs of the hind tibia and basitarsus are reduced, but not lost altogether. In most Megachilidae, however, the scopal hairs have moved entirely to the venter of the abdomen without retaining the ancestral scopae. I believe the most likely explanation is that the ancestor to Megachilidae evolved extensive scopal hairs that covered the legs and abdomen (similar to modern-day *Systropha*), and then the scopae was reduced for some unknown reason, leaving only the abdominal scopae. Some basal groups of Megachilidae, such as the genus *Apidosmia,* retain scopal hairs on the hind legs (Gonzalez et al. 2012) and may provide clues as to why other Megachilidae have apparently lost hind leg scopae.

Another open question is why some bee groups have not undergone significant scopal expansion despite transporting dry pollen. Examples of this include the genera *Anthophora* and *Xylocopa,* which transport surprisingly small pollen loads primarily on the hind tibia and basitarsis with only a little bit on the hind femur (Roberts and Vallespir 1978). One potential explanation is that they may supplement the external pollen loads with pollen transported internally in the crop. This is thought to occur in *Xylocopa* (Roubik 1989), but whether this also occurs in other groups like *Anthophora* is unknown.

#### The evolution of oil transport

The evolution of external transport of oil-moistened pollen is unclear and I have not had the opportunity to perform a firsthand investigation of oil-transporting bees. However, some level of inference can still be made about the evolution of this behavior based on known facts. Most importantly, oil-collecting bees also temporarily accumulate pollen on a specialized hair patch on the venter, just like bees that moisten the pollen with nectar. This pollen-accumulating patch appears to be much more extensive in oil-collecting bees; for example, the pollen-accumulating hairs in *Macropis* take up nearly the entire underside of the bee, including the abdominal sterna as well as the venter of the thorax (Cane et al. 1983, Schäffler and Dötterl 2011). The shared behavior of accumulating pollen on the venter suggests that oil-transport may have evolved from bees that originally moistened with nectar. This hypothesis is supported by the findings that the provisions of oil-collecting bees have been found to contain appreciable amounts of sugars. For example, the provisions *Centris maculifrons* contains glucose and fructose in addition to oil (Neff and Simpson 1981), and *C. brethesi* provisions also contained large amounts of sugars (Simpson et al. 1990). However, for these bees, it’s not clear if these sugars came from nectar transported in the crop and later added to provisions, versus whether the scopae also transport pollen moistened with some amount of nectar in addition to oil. More information is needed and the evolution of external transport of pollen moistened with oil remains an open question.

#### The secondary evolution of crop transport

Despite my hypothesis that internal pollen transport is the original form of pollen transport in bees, most if not all known examples of bees that transport pollen internally represent secondary evolutions of this behavior. There have been two instances where this has been examined in-depth: in the genus *Leioproctus* (Houston 1981) and in the *Colletes fasciatus-group* (Kuhlmann 2006). In both cases, the bees evolved from ancestors that transported dry pollen on external scopae (Houston 1981, Kuhlmann 2006). The switch to internal pollen transport is thought to be associated with a switch from hosts with large pollen to small pollen; this switch resulted in scopae that were ill-adapted to carry these fine pollen grains, driving the evolution to internal transport (Houston 1981, Kuhlmann 2006).

One intriguing possibility is that internal pollen transport has evolved repeatedly from dry transport because it never quite went away entirely. In other words, at least some bees that transport pollen externally may have never completely lost internal pollen transport and continued to transport a portion of pollen in the crop. Partial internal transport is something that has been mentioned by a couple of authorities on bee behavior and evolution, but to my knowledge it has never been investigated in depth. For example, Roubik (1989) states “A number of nonparasitic bees, for example *Ceratina* and *Xylocopa*, appear to collect pollen in both manners [internally and externally] and display a moderate reduction of scopae. Explanations for this behavior are lacking.” In addition, Michener (2007) cryptically states: “Finally, although pollen in bees’ crops is partly used for their own nutrition, some is carried to the nests and regurgitated.” If crop pollen transport was never completely lost, it would help explain why it has been able to evolve repeatedly in disparate bee lineages.

#### The secondary evolution of moist transport

The secondary evolution of moist transport from dry transport appears to be rare in bees. While there are some relatively well-documented examples of bees evolving dry or glazed transport from ancestors that transported moistened pollen (e.g. Portman and Tepedino 2017), there are no well-documented examples of the reverse (though it’s not clear how hard anyone has looked). This can be explained, at least in part, by the loss of behavior and structures to accumulate pollen on the venter. Most bees that transport dry pollen have lost the specialized patch of hairs to temporarily accumulate pollen and instead pass the pollen directly to the scopae or even gather it directly with the scopae. Without the temporary accumulation step, the behavior used to moisten the pollen before transferring it to the scopae is lost.

However, bees can potentially secondarily evolve moist transport if they develop a different behavior to moisten the pollen. This appears to be the case in at least some *Andrena* (*Dactylandrena*) species. For example, within the BBSL collection, there are specimens of *Andrena* (*Dactylandrena*) *porterae* that appear to have moistened pollen in the scopae. These bees gather pollen from the inaccessible flowers of *Ribes* using the mouthparts. The act of gathering pollen directly with the mouthparts can provide a mechanism to moisten the pollen with nectar that does not require passing the pollen up to the mouthparts to be moistened, as is done when bees temporarily accumulate pollen on the venter. However, there are also many bees that gather pollen with the mouthparts but still clearly transport dry pollen, so the steps driving the secondary evolution of moist pollen transport are not entirely clear and require more investigation.

### Parallel evolution in pollen wasps

In addition to bees, an evolutionary change to provisioning the young with pollen from an ancestral predatory lifestyle has arisen in two other hymenopteran lineages. These examples can inform about how this process occurred in bees. The two examples include the masarid pollen wasps in Vespoidea, and the genus *Krombeinalictus* in Crabronidae. The biology of the single species of *Krombeinalictus* is poorly known, so the lessons that can be learned from it are limited (Krombein and Norden 1997). However, the biology masarid wasps are relatively well-known, and offer a valuable source of information regarding the evolution of pollen provisioning from an ancestral predatory lifestyle.

Using masarid pollen wasps (hereafter referred to as “pollen wasps”) as a template, we can compare them to the proposed sequence of bee evolution. This is important because there are many pollen wasps that have a life history similar to the hypothesized protobee. This demonstrates that the proposed stages of bee evolution are not just abstract intellectual constructs, but instead represent viable life-history strategies that exist in the present day.

#### Hypothesis: Crop transport is ancestral and it evolved from ancestral adult pollen-feeding behavior

All known pollen wasps transport pollen internally, making it clear that it is the ancestral form of pollen transport. Similar to what I have hypothesized for bees, internal transport in pollen wasps is thought to have evolved from ancestral pollen feeding behavior, in this case in stem-group vespid wasps that consumed pollen as adults but provisioned their larvae with prey (Mauss 2007). The antiquity of pollen-feeding behavior in adults is further supported by the ubiquity of this behavior in the present day, where adult pollen wasps of both sexes consume pollen for their own nutritional needs (Mauss et al. 2005, 2019). This is most well-documented in males, of which multiple species have been observed collecting pollen and dissections have found pollen in their crop (Mauss and Müller 2000, 2016, Mauss et al. 2003, 2005, 2006, Groddeck et al. 2004). Because females transport pollen internally, it’s difficult to determine whether the pollen they consume is for provisions or their own nutrition. However, dissection of female *Pseudomasaris edwardsii* revealed pollen in the mid- and hindgut, confirming that they consumed pollen for their own nutrition (Torchio 1970). These examples suggest that pollen consumption is widespread in adult pollen wasps.

#### A discussion of the mechanisms by which pollen wasps feed on pollen

Like bees, pollen wasps gather pollen in two ways, nibbling directly with the mouthparts and by drawing the foreleg through the mouthparts. Nibbling pollen directly with the mouthparts is present in many pollen wasps (Mauss et al. 2019) and likely represents the ancestral form. This type of pollen gathering is most well-documented in *Pseudomasaris edwardsii* (Torchio 1970, Neff and Hook 2007), *Quartinia tenerifinia* (Mauss and Mauss 2016), and *Ceramius hispanicus* (Krenn et al. 2002). As in bees, nibbling directly with the mouthparts appears to be relatively rare and drawing the forelegs through the mouthparts to consume pollen is the more common form. Indeed, a pollen-comb on the galea has been found in pollen wasps, where it is presumably used to remove pollen from the forelegs as they are drawn through the mouthparts (Krenn et al. 2002, Mauss et al. 2019). Multiple species of pollen wasp have been documented to gather pollen through a combination of nibbling with the mandibles and drawing the forelegs through the mouthparts. This is seen in species such as *Celonites fischeri* (Mauss and Müller 2014), *Ceramius fonscolombei* (Mauss et al. 2003), *Quartinia canariensis* (Mauss and Müller 2016), and *Quartinia major* (Mauss et al. 2018). The use of forelegs in pollen gathering may be related to the accessibility of the pollen; *C. hispanicus* is reported to nibble pollen when anthers are accessible, and uses the forelegs when they are not (Mauss and Müller 2000, Krenn et al. 2002).

#### Temporary accumulation of pollen in pollen wasps

Similar to bees, many species of pollen wasps also temporarily accumulate pollen, with the pollen initially gathered onto a specialized patch of hairs before being brought to the mouthparts by the forelegs (Müller 1996). The most well-documented examples of the temporary accumulation of pollen in wasps include species that first accumulate pollen on the face, often on knobbed or hooked hairs (Müller 1996, Mauss 2006, Mauss et al. 2016). Other pollen wasps accumulate pollen on the dorsum of the thorax via “rasping” behavior (Torchio 1974, Portman et al. 2019). Most importantly, there are pollen wasp species that gather pollen by first accumulating pollen on the venter of the thorax. For example, *Rolandia maculata* has a specialized patch of stiff hairs with bent tips on the venter of the thorax; this patch accumulates pollen before being ingested using the forelegs (Houston 1995). A similar pollen-accumulating hair patch is found on the venter of *Ceramius braunsi* (Gess and Gess 1989). Although Gess and Gess (1989) describe the pollen gathering in *C. braunsi* as being performed solely by the forelegs, without an accumulation step, the accumulation of pollen in the ventral hair patches suggests Gess and Gess (1989) may have missed that behavior.

#### Tying back to bees

Although there are no pollen wasps that are known to transport pollen externally, there are still important parallels to the hypothesized evolution of pollen transport in bees. Specifically, in both bees and pollen wasps, adults feed on pollen for their own nutritional needs and they can consume pollen either through nibbling or drawing the foreleg through the mouthparts. Importantly, crop transport of pollen is unambiguously ancestral in pollen wasps, and some pollen wasps share the behavior of temporarily accumulating pollen on the venter. It is especially striking that there are pollen wasps that gather and transport pollen the same way that *Perdita tortifoliae* gathers and consumes pollen, which lends credence to the hypothesis that temporarily accumulating pollen on the venter (as exemplified by *Perdita tortifoliae* in earlier sections) represents an ancestral form of pollen transport in bees. However, masarids have clearly never made the evolutionary transition to external transport. The lack of this evolutionary innovation could help explain why bees are so much more diverse than masarids, despite their similar evolutionary ages. Overall, this supports the hypothesis that bees and masarids followed a similar evolutionary pathway in the initial stages of the evolution of pollen transport.

## Conclusion

In this paper I have laid out a hypothesis on the origin and evolution of pollen transport in bees. Under this view, internal transport in the crop represents the original pollen transport behavior and it evolved from pollen feeding in adults. From there, bees evolved the ability to temporarily accumulate pollen on a specialized patch on the venter of the thorax, which represents a necessary transition stage that led to external transport of pollen moistened with nectar on the hind legs. External transport of dry or glazed pollen then evolved from external moist transport. Finally, the evolution of external dry pollen transport led to the expansion of the scopal hairs in many bee groups. This hypothesis is supported by multiple lines of evidence, particularly by observations on present-day pollen-feeding and pollen-gathering behavior in bees which allow us to reconstruct the evolutionary history of these behaviors. Importantly, comparing the evolution of pollen transport of bees and pollen wasps boosts this hypothesis because it highlights potential paths of parallel evolution and demonstrates that the hypothesized transition forms in bees are actually viable life history strategies that exist in the present day in some pollen wasps.

Under the hypothesis laid out here, the evolution of external pollen transport in bees can be reconstructed by examining the steps of present-day pollen gathering behavior. In the present day, the transport of moistened pollen requires a transition step (temporary accumulation of pollen on the venter) that results in pollen taking a complicated and circuitous route: first the pollen is picked up by the forelegs, then transferred to a temporary holding area on the venter of the thorax, this pollen is then picked back up by the forelegs, brought forward to the mouthparts where it is moistened with nectar, passed backwards again where it is scraped off the forelegs by the midlegs before finally being deposited onto the hind legs. However, this process can be explained if it is viewed as the result of external moist pollen transport evolving by simply adding additional behaviors onto internal pollen transport; in moist transport, the original behavior of bringing pollen forward to be consumed by the mouthparts is retained, but instead of being consumed, the pollen is instead moistened and passed back the hindlegs. Most importantly, each individual stage of this evolutionary process is adaptive in its own right. The consumption of pollen via the foreleg and the temporary accumulation of pollen are both behaviors that are seen in the present day in both bees and pollen wasps.

My hypothesis that internal pollen transport is ancestral in bees marks a return to the earliest hypotheses regarding the genesis of bees, which was first laid out by Müller (1883) and expanded by Malyshev (1969) and Jander (1976). All of the previous workers cited the hairless bodies, poorly-developed pollen brushes, short tongues, and similarity to sphecid wasps as evidence that *Hylaeus* represented an ancestral bee group. Although recent molecular phylogenies have made it clear that *Hylaeus* and other Colletidae are not basal (Danforth et al.

2012), it does not negate the fact that the protobee almost certainly did have many of those characteristics, particularly poorly-developed body hairs and pollen-collecting structures and behaviors. In other words, the fact that *Hylaeus* are not basal does not invalidate the other logical arguments in favor of crop transport being ancestral. In particular, the parallel evolution with pollen wasps is one of the strongest arguments in favor of crop transport being ancestral, which is further bolstered by the degree of similarity in their evolutionary development laid out in the previous section.

Under this framework, I contend that moist pollen transport is ancestral to dry pollen transport. This represents the first detailed hypothesis about how moist pollen transport could have evolved (with perhaps the exception of Michener et al. (1978)), and it marks a deviation from the conventional wisdom that moist transport evolved from dry transport (Müller 1883, Michener 1944, Michener et al. 1978, Roberts and Vallespir 1979, Pasteels et al. 1983). The assumption that moist transport is the more derived character seems to stem, at least in part, by the notion that Apidae, and especially honeybees, represent the most advanced or “most derived” bees (e.g. Müller 1883, Jander 1976, Michener 1979). The hypothesis that moist transport represents the ancestral form of external pollen transport makes sense because it does not require specialized morphological characters such as well-developed branched hairs or scopae. Indeed, it allows pollen types of a wide variety of sizes and shapes to be carried on short and sparse simple hairs instead of the scopal adaptations typically seen in bees that transport dry pollen (Roberts and Vallespir 1978, Portman and Tepedino 2017, Danforth et al. 2019). In contrast, the evolution of dry transport from moist transport is associated with the elaboration, specialization, and expansion of the scopal hairs (Portman and Tepedino 2017).

Most importantly, the hypotheses laid out here create a consistent framework that is informed by present-day bee behavior and allows us to make broad predictions about the biology and evolution of bees. The most important of these predictions are laid out below:

1. Additional studies on Melittidae and other basal bees will reveal that most groups transport moistened pollen.
2. Most bees that transport moistened pollen temporarily accumulate pollen on the venter (or gather pollen directly with mouthparts). The obvious exceptions here are *Apis* and *Bombus* (but not *Trigona s.l.*, see Michener et al 1978); it’s not clear why this is the case but this should be a derived condition.
3. Additional studies will also reveal that species that gather pollen by accumulating pollen on a patch of specialized hairs on the venter also accumulate on that patch when feeding on pollen.
4. In bee lineages where there has been a transition between moist and dry external transport, moist transport will be found to be ancestral (except when pollen is gathered directly with the mouthparts).
5. The evolutionary transition of moist transport to dry transport will be associated with the use of pollen that is particularly adhesive, large, or spiny, which would make them more efficiently transported dry (e.g. Portman and Tepedino 2017).
6. Investigation of bees that transport pollen externally will reveal bees that transport a portion of pollen internally as well. This is particularly relevant for Melittidae and bees with small scopae.
7. Additional studies of the pollen gathering behavior of pollen wasps will reveal species that gather pollen by temporarily accumulating it on the venter before transferring to the mouthparts (as in Houston 1995).
8. The broad and flattened hind basitarsus, a character shared by all bees that separates them from wasps (Radchenko and Pasenko 1996, Engel 2001, Michener 2007), is a result of that being the location of the original external scopae. This suggests that the most recent common ancestor of all extant bees transported external moist pollen on the hind legs.
9. Evolutionary trends will reveal that bees have undergone an expansion of the area of scopal hairs from the ancestral location on the hind tibia and basitarsus (rather than the reverse — a consolidation onto the hind tibia and basitarsus).

The last prediction stands in strong contrast to the primary competing hypothesis regarding the origin of pollen transport, originally proposed by Radchenko and Pasenko (1996) and supported by Michener (2007). Under that hypothesis, external dry transport is ancestral and scopal hairs coalesced and specialized from a diffuse and unspecialized ancestral form. This creates a key difference between their hypothesis and my own. Under my hypothesis, where moist transport is ancestral to dry transport, the hind tibia and basitarsus are the ancestral location of the scopa, and all modern-day external scopae have expanded outward from there. In contrast, under the hypothesis of Radchenko and Pasenko (1996), the reverse would be predicted — that diffuse scopae should coalesce on the hind tibia and basitarsus. My prediction that scopae that transport dry pollen will have become increasingly proximal rather than increasingly distal also stands in contrast to the conventional wisdom regarding the evolution of pollen transport (Pasteels and Pasteels 1979, Thorp 1979, Pasteels et al. 1983, Westerkamp 1996). Based on the evidence currently available (e.g. Roberts and Vallespir 1978), expansion of the scopae, rather than the consolidation, appears to be the rule, though this has yet to be rigorously tested from a phylogenetic standpoint.

Here, I have presented the first detailed hypothesis of how external moist transport could have evolved and this marks a step forward in a field that has seen little progress despite the major advances in our understanding of bee phylogenies and deep evolutionary relationships. Further, this framework allows for us to better understand bee biology in the present day, and offers an evolutionary explanation for behaviors, such as the temporary accumulation of pollen on the venter, that may at first seem incongruous. It is my hope that this will stimulate the research needed to confirm or refute this hypothesis. While better-resolved phylogenies would certainly be helpful, answers about the origin and evolution of pollen transport primarily require studies on the natural history, behavior, and functional morphology of bees and related Hymenoptera.

## Acknowledgments

I thank Vince Tepedino for feedback on various drafts of this manuscript. I acknowledge the support from the Microscopy Core Facility at Utah State University for the SEM work. The fieldwork that led to most of the bee observations that form the basis of this work was made possible due to funding by the U.S. Fish and Wildlife Service (Grant F16AP00680).

